# On the role of arkypallidal and prototypical neurons for phase transitions in the external pallidum

**DOI:** 10.1101/2021.01.06.425526

**Authors:** Richard Gast, Ruxue Gong, Helmut Schmidt, Hil G.E. Meijer, Thomas R. Knösche

## Abstract

The external pallidum (GPe) plays a central role for basal ganglia functions and dynamics and, consequently, has been included in most computational studies of the basal ganglia. These studies considered the GPe as a homogeneous neural population. However, experimental studies have shown that the GPe contains at least two distinct cell types (prototypical and arkypallidal cells). In this work, we provide in silico insight into how pallidal heterogeneity modulates dynamic regimes inside the GPe and how they affect the GPe response to oscillatory input.

We derive a mean-field model of the GPe system from a microscopic spiking neural network of recurrently coupled prototypical and arkypallidal neurons. Using bifurcation analysis, we examine the influence of the intra-pallidal connectivity on the GPe dynamics. We find that under healthy conditions, the inhibitory coupling determines whether the GPe is close to either a bi-stable or an oscillatory regime. Furthermore, we show that oscillatory input to the GPe, arriving from subthalamic nucleus or striatum, leads to characteristic patterns of cross-frequency coupling observed at the GPe. Based on these findings, we propose two different hypotheses of how dopamine depletion at the GPe may lead to phase-amplitude coupling between the parkinsonian beta rhythm and a GPe-intrinsic gamma rhythm. Finally, we show that these findings generalize to realistic spiking neural networks of sparsely coupled type-I excitable GPe neurons.

**Significant Statement:** Our work provides (a) insight into the theoretical implications of a dichotomous GPe organization for its macroscopic dynamic regimes, and (b) an exact mean-field model that allows for future investigations of the relationship between GPe spiking activity and local field potential fluctuations. We identify the major phase transitions that the GPe can undergo when subject to static or periodic input and link these phase transitions to the emergence of synchronized oscillations and cross-frequency coupling in the basal ganglia. Due to the close links between our model and experimental findings on the structure and dynamics of prototypical and arkypallidal cells, our results can be used to guide both experimental and computational studies on the role of the GPe for basal ganglia dynamics in health and disease.

## Introduction

The basal ganglia (BG) are a set of interconnected subcortical nuclei that form different feedback loops with cortex and thalamus (Alexander and Crutcher, 1990; Bolam et al., 2000). Due to its recurrent connections with nearly all other BG nuclei, the globus pallidus pars externa (GPe) plays a major role for information transmission through the BG (Kita, 2007). In Parkinson’s disease (PD), synchronized oscillations have been reported throughout all major BG nuclei (Wichmann, 2019) including the GPe (Wichmann and Soares, 2006; Mallet et al., 2008). These oscillations are characterized by transient power increases in the beta frequency band (12-30 Hz) and an increased phase-amplitude coupling between the phase of a beta signal and the amplitude of a high-frequency gamma signal (50-250 Hz) (Jenkinson et al., 2013; Lofredi et al., 2019; Gong et al., 2020). Computational models of BG phase transitions in PD suggest that the GPe is involved in the oscillation generation, either via its recurrent coupling with the subthalamic nucleus (STN) or via its processing of inputs from striatum (STR) (Pavlides et al., 2015; Schroll and Hamker, 2016; Rubin, 2017). Most of these computational models regarded the GPe as a homogeneous population of neurons. However, two major cell types have been identified within the GPe, which differ in their electrophysiological properties, firing rates, and firing patterns: Prototypical (GPe-p) and arkypallidal (GPe-a) cells (Cooper and Stanford, 2000; Abdi et al., 2015; Hegeman et al., 2016). Regarding their efferent synapses, it has been shown that GPe-p cells preferentially project to STN and BG output nuclei, whereas GPe-a cells provide feedback to STR (Mallet et al., 2012; Hernández et al., 2015). Furthermore, a recent study found STN and STR to differentially affect GPe-p and GPe-a in mice (Ketzef and Silberberg, 2020). Regarding cell-type specific differences in GPe-intrinsic axon collaterals, there is evidence from mice experiments that prototypical cells express more numerous axon collaterals than arkypallidal cells (Mallet et al., 2012; Ketzef and Silberberg, 2020). Still, a substantial amount of arkypallidal axon collaterals was identified that targeted prototypical GPe cells (Mallet et al., 2012).

Depending on the pattern of mutual inhibition between those two major GPe cell populations, different modes of GPe internal dynamics may exist. Asymmetric connections between the two cell types may give rise to a feed-forward inhibition scenario, where an excitatory input to one population could silence the other population. Alternatively, winner-takes-all (WTA) dynamics can arise in scenarios of mutual inhibition be-tween two populations (Schmidt et al., 2018). Such regimes could be exploited by asymmetric inputs from STN and STR to GPe-a and GPe-p, which would allow for transient switching between the two different output pathways of the GPe. Therefore, we argue that the relationship between synaptic coupling and neural dynamics in a GPe composed of arkypallidal and prototypical cells could be a major factor to understand information routing in the BG. In this study, we examine the effects of different GPe coupling patterns on GPe behavior. For this purpose, we derive and analyze a mean-field description of two fully coupled inhibitory populations, following the approach by (Luke et al., 2013; Montbrió et al., 2015). Importantly, this mean-field description captures the exact macroscopic dynamics of the underlying, heterogeneous spiking neural network and can thus capture population-intrinsic spike resonance phenomena that classic mean-field approaches would miss. This in itself makes our modeling approach interesting for the understanding of synchronization processes inside the GPe. In an initial analysis of the two-population GPe model, we identify mono-stable, bi-stable and oscillatory regimes via bifurcation analysis, the existence of which depends on the GPe-intrinsic coupling pattern. We then show that the GPe expresses distinct responses to periodic input when initialized in either of these regimes. Finally, we analyze how the macroscopic phase transitions found in the GPe mean-field model translate to spiking neural networks with realistic numbers of neurons and axons.

## Materials and Methods

### Model Definition

#### Mathematical Formulation of Population Dynamics

We consider the GPe as a nucleus of two distinct populations of GABAergic projection neurons (Kita, 2007; Hegeman et al., 2016). Both populations, prototypical and arkypallidal neurons, express high average spontaneous firing rates of 50-70 Hz and 20-40 Hz, respectively (DeLong, 1971; Cooper and Stanford, 2000; Wichmann and Soares, 2006; Jaeger and Kita, 2011). To model synaptic influences on the spike timings of GPe neurons, it is important to know their type of excitability. This can be inferred from their phase-response curve (Gutkin et al., 2005). Neurons can either express type-I excitability, meaning that the direction in which the excitability of a neuron is changed by extrinsic input is not dependent on the intrinsic phase of the neuron, or express type-II excitability, meaning that the direction in which the excitability of a neuron is changed by extrinsic input does depend on the intrinsic phase of the neuron (Izhikevich, 2000). While computational studies demonstrated that both type-I and type-II excitability can be identified in single cell models of GPe neurons (Schultheiss et al., 2010; Fujita et al., 2012), experimental investigations only revealed type-I excitability so far (Wilson, 2013). Furthermore, it has been shown that coupled networks of type-I excitable neurons can express type-II excitability on the network level (Dumont and Gutkin, 2019). Thus, as a base neuron model, we use the quadratic integrate-and-fire neuron (QIF), which is the canonical form of type-I excitable neurons and expresses a quadratic and thus non-linear input-output relationship (Izhikevich, 2000). This choice also accounts for the non-linear input-output relationship reported in prototypical and arkypallidal cells (Kita, 2007; Schultheiss et al., 2010; Fujita et al., 2012; Abdi et al., 2015). The evolution equation of the *j^th^* QIF neuron embedded within either the GPe-p or GPe-a is given by

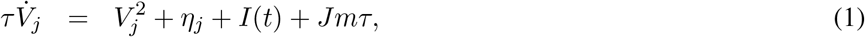

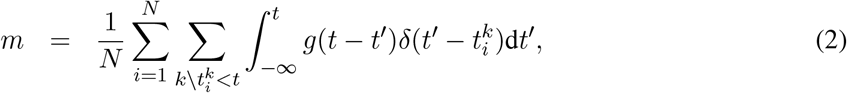

with neural excitability *η_j_*, synaptic strength *J*, evolution time constant *τ*, extrinsic input *I*(*t*) and synaptic activation *m*. A neuron *j* generates its *k^th^* spike at time 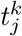 At this time, it reaches the spiking threshold *V_θ_* and the membrane potential *V_j_* is reset to a reset potential *V_r_*. The integral kernel *g*(*t* − *t*′) represents synaptic dynamics, e.g. in the case of mono-exponential synapses 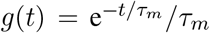 with synaptic time scale *τ_m_*. We introduce the exact shape and timescales of *g* in the following sub-section. Equations (1) and (2) represent an all-to-all coupled network of *N* QIF neurons with homogeneous connection strengths *J*. Assuming all-to-all connectivity as well as infinitely large neural populations, we can use the mean-field model proposed in (Montbrió et al., 2015). The authors derived a set of two coupled differential equations describing the evolution of the macroscopic firing rate *r* and membrane potential *v* of the QIF population given by (1) and (2):

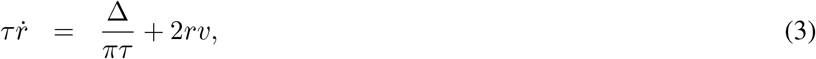

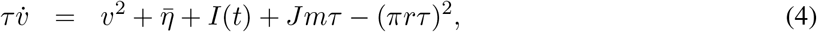

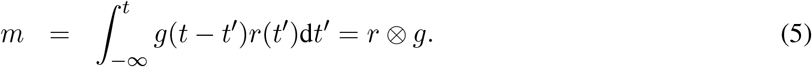

Here, the synaptic activation *m* takes the form of a simple convolution of the average firing rate *r* with the synaptic response kernel *g*, henceforth abbreviated by the convolution operator ⊗. The parameters 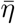 and Δ are the center and half width at half maximum of a Lorentzian distribution over the single neuron parameters *η_j_*. Thus, 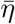 and Δ allow to control the average and heterogeneity of the firing rates inside the QIF population, respectively. Spontaneous firing rates of GPe cells cannot be explained by glutamatergic input alone, since brain slice recordings still showed autonomous activity of up to 26 Hz after synaptic transmission was blocked pharmacologically (Günay et al., 2008). In other words, GPe cells are strong pacemaker cells that show regular firing at a cell-specific frequency under synaptic isolation (Mercer et al., 2007). Across GPe cells, a substantial amount of heterogeneity of the intrinsic firing frequencies has been reported (Wilson, 2013). By considering the background excitabilities *η_j_* as distributed quantities, we account for these findings.

We are aware that the all-to-all coupling and infinite population sizes are in contrast to the actual GPe structure (Wilson, 2013; Hegeman et al., 2016). However, it has been recently shown that the mean-field model predictions can generalize to a fairly wide range of network sizes and coupling probabilities (Gast et al., 2020). Even for QIF networks with recurrent coupling probabilities of 1%, the authors found that population sizes of *N* = 8000 neurons were sufficient to accurately reproduce the macroscopic dynamics predicted by the mean-field model. Given that population sizes of primate GPe are on the order of 10^5^ and recurrent coupling probabilities are around 5% (Wilson, 2013), we expect that this mean-field model is sufficient to capture the macroscopic dynamics of QIF populations with realistic cell counts and coupling probabilities.

#### Mathematical Formulation of Axonal Propgation and Synaptic Dynamics

In a next step, we define the coupling function *g* which, in our model, acts as a lumped representation of axonal propagation and synaptodendritic integration. In other words, *g* serves to link single spikes emitted by neuron *j* to changes in the membrane potential of any other neuron. GPe to GPe connections have been suggested to express axonal transmission delays of around 1.0 ms (Jaeger and Kita, 2011) and make use of GABAergic synapses (Kita, 2007). Since axon collaterals can express a substantial variability in individual axon diameters and myelination properties (Schmidt and Knösche, 2019), we modeled the axonal transmission delays via gamma distributions (Smith, 2011). The probability density function of the gamma distribution can be written as

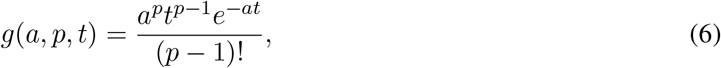

with shape parameter *p* and scale parameter *a*. These parameters can be used to control the mean *μ* and width *σ* of the delay distribution via the functional relationships 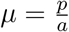 and 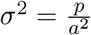 (Smith, 2011). Choosing (6) as functional form of the function *g* in equation (5), the synaptic convolution operation can be approximated by the following set of coupled ordinary differential equations (ODEs):

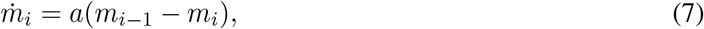

where *i* = 0, 1, 2,…, *p* and *m*_0_ = *r* (Smith, 2011). Using this formulation, the number of coupled ODEs depends on the shape parameter of the gamma function, which means that the overall dimensionality of the system depends on the order parameters *p* at each synaptic connection in the model.

In addition to the axonal delays, we also included a dynamic model of the electrochemical processes that lead to a change in the post-synaptic potential after a pre-synaptic action potential traveled down the axon. A popular choice to express these dynamics is via a convolution with a bi-exponential synaptic response kernel, for which the rise and decay time constants are specific to the type of pre- and post-synapse (Deco et al., 2008). Such a bi-exponential synaptic response function is given by

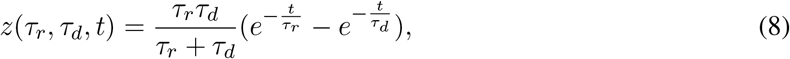

with *τ_r_* and *τ_d_*, denoting the synaptic rise and decay time constants, respectively. A convolution of the delayed axonal response *m_p_* with (8) can be approximated by two coupled ODEs of the form:

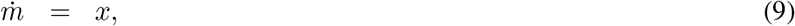

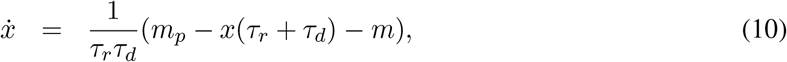

with *m* being the final synaptic input entering into (3). Thus, we specify the convolution integral expressed by (5) in our model as subsequent convolutions of *r* with the gamma function (6) and the bi-exponential function (8), allowing us to capture the characteristics of both axonal delay distribution and post-synaptic currents.

#### Specification of the two-population GPe Model

Based on these dynamic equations for neural populations and synaptic transmission, we can now introduce the full set of equations of our GPe model. Since the number of equations of the ODE approximation (7) to the gamma kernel convolution (5) depends on the parameter *p* of (6), we chose to provide a set of integro-differential equations for generality and brevity. However, for our results, each gamma kernel convolution was formulated as a set of coupled ODEs of the form (7) and each convolution with a synaptic response kernel of the form (8) was formulated as the ODE system given by (9) and (10). The following set of coupled integro-differential equations describes the average firing rate and average membrane potential dynamics at GPe-p and GPe-a:

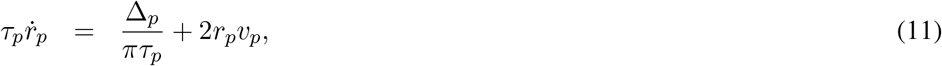

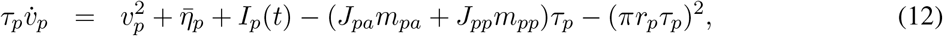

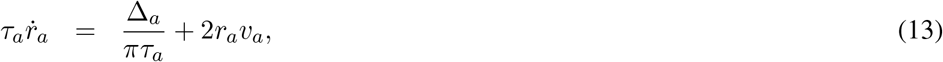

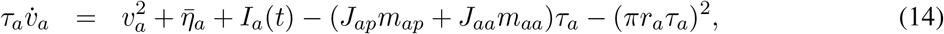

where *p* and *a* are the subscripts for prototypical and arkypallidal GPe, respectively, and subscripts of the form *A_xy_* represent the variable *A* that is specific to the synaptic transmission from population *y* to population *x*. Hence, each synaptic response function *g_xy_* is specific to a given synaptic transmission and takes the form

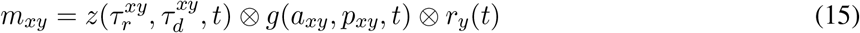

with connection specific synaptic rise and decay times 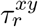 and 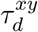 and connection specific axonal delay distribution shape and scaling *a_xy_* and *p_xy_*.

#### Mathematical Formulation of Extrinsic Model Inputs

Extrinsic input can generally be applied via the extrinsic forcing parameters *I_p_*(*t*) and *I_a_*(*t*) to GPe-p and GPe-a, respectively. In our simulations, we applied step function inputs to each of the populations. These are defined as

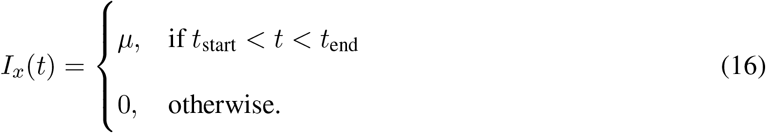

Here, *μ* defines the input strength, whereas *t* _start_ and *t* _end_ define the beginning and end of the time interval in which the input is applied. Furthermore, we also applied periodic input to the GPe-a. We used the Stuart-Landau oscillator as the generating model of a sinusoidal signal with period *ω* (Fujimura, 1997):

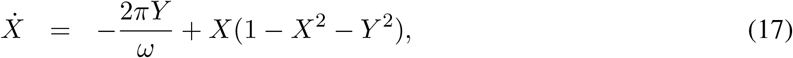

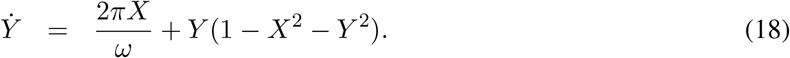

Additionally, to account for the bursting characteristics of typical striatal inputs arriving at the GPe (Jaeger et al., 1995), we applied a sigmoidal transformation to the Stuart-Landau oscillator, giving us the final input

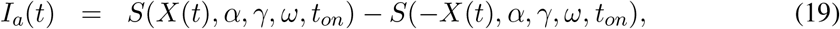

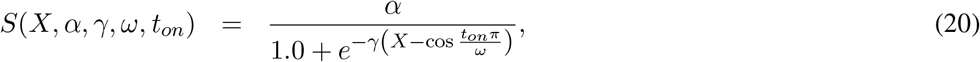

where *S* represents a sigmoidal transform with maximum *α* and steepness *γ*. The cosine term ensures that the input *I_a_*(*t*) expresses bursts around the maxima and minima of *X*. We set the steepness of the bursts to *γ* = 100.0 and the width of the bursts to *t_on_* = 5.0ms. For a more detailed description of this sigmoidal transformation of a sinusoidal signal, see (Lourens et al., 2015).

### Model Parameters

The dynamics at GPe-p and GPe-a are each governed by membrane time constants *τ* and two parameters 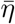 and Δ that determine the center and half width at half maximum of the distribution of single cell firing rates inside the populations. Additionally, the four synaptic connections between GPe-p and GPe-a are each parameterized via a lumped synaptic strength *J*, two axonal delay parameters *μ* and *σ* and the synaptic rise and decay time constants *τ_r_* and *τ_d_*. To find a parameterization of the model that resembles realistic macroscopic neural dynamics inside the GPe, all time constants in the model were informed by experimental data. The values of those parameters and their sources are listed in Table 1. The rest of the parameters are either scaling parameters of synaptic strengths or direct input currents to the populations and were chosen such that the steady-state firing rates of the model reflect average firing rates recorded in the GPe of healthy monkeys (Kita et al., 2004; Wichmann and Soares, 2006). GPe coupling patterns are defined by the four coupling strengths *J_pp_*, *J_pa_*, *J_ap_*, *J_aa_*. We re-defined these coupling strengths for a more systematic investigation of GPe coupling patterns: 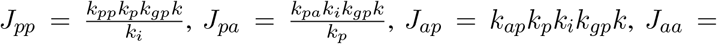 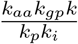.Thus, *k_p_* represents the relative strength of GPe-p projections over GPe-a projections, *k_i_* represents the relative strength of projections between GPe-p and GPe-a as compared to projections within GPe-p and GPe-a, and *k_gp_* represents the scaling of all GPe-to-GPe connections. If not defined otherwise, these parameters were set to the default values reported in Table 1.

**Table 1:** Model Parameters

| Parameter | Value | Reference | Parameter | Value |
| --- | --- | --- | --- | --- |
| $\tau_p$ | 25.0 ms | Cooper and Stanford (2000) | $\Delta_p$ | 90.0 |
| $\tau_a$ | 20.0 ms | Cooper and Stanford (2000) | $\Delta_a$ | 120.0 |
| $\mu_{pp}, \mu_{aa}, \mu_{pa}, \mu_{ap}$ | 1.6 ms | Jaeger and Kita (2011) | $\eta_p$ | $\frac{10}{3}\Delta_p$ |
| $\sigma_{pp}, \sigma_{aa}, \sigma_{pa}, \sigma_{ap}$ | 0.4 ms | Jaeger and Kita (2011) | $\eta_a$ | $\frac{5}{6}\Delta_a$ |
| $\tau_r^{pp}, \tau_r^{aa}, \tau_r^{pa}, \tau_r^{ap}$ | 0.5 ms | Sims et al. (2008) | $k_{gp}$ | 300.0 |
| $\tau_d^{pp}, \tau_d^{aa}, \tau_d^{pa}, \tau_d^{ap}$ | 5.0 ms | Sims et al. (2008) | $k_p$ | 1.5 |
| - | - | - | $k_i, k_{pp}, k_{aa}, k_{pa}, k_{ap}$ | 1.0 |

### Model Analysis

To analyze the behavior of the model given by (11–14), we employed the open-source Python toolbox PyRates (Gast et al., 2019). We chose PyRates’ interface to the SciPy Runge-Kutta solver with adaptive integration step-size (Virtanen et al., 2020) for numerical integration of the model dynamics for a given initial condition. For bifurcation analysis, we used PyRates’ interface to Auto-07p (Doedel et al., 2007) which allows to perform numerical parameter continuation and automatic bifurcation detection. For an in-depth explanation of these techniques see (Meijer et al., 2009; Kuznetsov, 2004), for example. To analyze the behavior of the spiking neural networks corresponding to our mean-field models, we employed custom Matlab code. Numerical integration of the spiking neural network dynamics was performed via an explicit Euler algorithm with an integration step-size of 0.001 ms, which we found to be sufficiently small to capture all model dynamics. The scripts and configuration files for all simulations and parameter continuations are available at the following public Github repository: https://github.com/Richert/GPe_Dynamics.

### Spectral Analysis

We also analyzed the GPe model behavior in the frequency domain. To this end, we used time series of 320 seconds of simulated GPe-p firing rate dynamics sampled at 1 ms and cut off the first 20 s to remove initial transients from the time series. Power spectral densities (PSDs) were calculated from the raw simulation data using Welch’s method. We used FFT segments of length 2048 and an overlap between segments of 1024 time steps. For quantification of phase-amplitude coupling (PAC) and phase-phase coupling (PPC) between different frequency components of the GPe-p firing rate dynamics, we followed the procedure described in (Gong et al., 2020). PAC measures the amount of modulation of the amplitude of a high-frequency signal by the phase of a low-frequency signal and was evaluated by means of the Kullback-Leibler-based modulation index (KL-MI) (Tort et al., 2010). Both the low- and high-frequency signals were acquired by band-pass filtering the GPe-p firing rate time series. Following the procedure described in (Gong et al., 2020), we evaluated the KL-MI for multiple pairs of phases at frequencies *f_p_* ∈ 2, 4, 6,…, 30 Hz and amplitudes at frequencies *f_a_* ∈ 50, 60, 70,…, 250 Hz. For each pair of *f_p_* and *f_a_*, we filtered the GPe-p firing rate using an FIR band-pass filter centered at *f_p_* with a band-width of 2 Hz and using another FIR band-pass centered at *f_a_* with a band-width of *f_p_* Hz. We then applied the Hilbert transform to the two band-pass filtered signals and extracted the phase from the signal filtered around *f _p_* and the amplitude of the signal filtered around *f_a_*. Phases were then sorted into 16 bins and the amplitudes corresponding to each bin were averaged. Then, the KL-MI of the distribution of the average amplitude across phase bins was calculated as described in (Tort et al., 2010), which measures the difference to a uniform distribution. Furthermore, we evaluated PPC for the GPe-p firing rates filtered around *f_p_* and *f_a_* using the waveform analysis described in (Gong et al., 2020). In short, this method calculates the average waveform of the high-frequency signal, time-locked to the zero-crossing of the low-frequency signal. The resulting metric is bounded between 0 and 1, with *PPC* = 1 indicating that the phase of the high-frequency signal (filtered at *f_a_*) is always the same at zero-crossings of the phase of the low-frequency signal (filtered at *f_p_*). Hence, for a given GPe-p firing rate time series, we acquired a 15 × 21 PAC (PPC) matrix *C_pa_* (*C_pp_*) with entries for each pair of *f_a_* and *f_p_*. To evaluate the overall amount of PAC in a time series, we calculated the average across the PAC matrix (*mean PAC* in Fig. 3). To evaluate the similarity between PAC and PPC across low- and high-frequency components of a time series, we calculated the Pearson correlation coefficient between the PAC and the PPC matrix (*correlation(PAC,PPC)* in Fig. 3). Finally, to examine whether high PAC coincided with high or low PPC across pairs of *f_p_* and *f_a_* in a time series, we evaluated the average of *C_pa_* ∗ *C_pp_* or *C_pa_* ∗ (1 − *C_pp_*) (*mean PAC * PPC* and *mean PAC * (1-PPC)* in Fig. 3). Here, ∗ denotes element-wise multiplication of matrices.

## Results

In this section, we report the results of our analysis of the relationship between model parameters and neural dynamics for the GPe model given by (11–14). We focus on parameters that contribute to a difference between GPe-p and GPe-a, which include the coupling strengths within the GPe as well as additional inputs to the two populations.

### Effects of GPe-Intrinsic Coupling

As the first part of our analysis, we performed bifurcation analysis of the GPe mean-field model given by (11–14) to investigate whether different coupling patterns between prototypical (GPe-p) and arkypallidal (GPe-a) cells promote different macroscopic states and phase transitions. To this end, we defined different intra-pallidal coupling patterns, for each of which we performed parameter continuations in the input parameter *η_a_*. We did not consider scenarios in which the total GPe-a projection strength is stronger than the total GPe-p projection strength, i.e. *k_p_* < 1.0, since such coupling patterns seem unlikely given the GPe axon collaterals reported in (Mallet et al., 2012).

#### Coupling Pattern 1: Strong Coupling Between Prototypical and Arkypallidal Cells Promotes Bi-Stable GPe Regimes

We started out by investigating the case *k_i_* = 2.2 and *k_p_* = 1.0, i.e. a GPe coupling pattern with stronger coupling between than within GPe-p and GPe-a and equal total projection strengths of GPe-p and GPe-a (see Figure 1A). As can be seen in Figure 1C, we found two fold bifurcations when performing a single parameter continuation in *η_a_*. Changes to both *η_a_* and *I_a_*(*t*) can be considered to reflect changes in the average input to GPe-a neurons. Therefore, the fold bifurcations represent the outer boundaries of a bi-stable regime, in which transient inputs to GPe-a (or GPe-p) allow to switch between two stable states (see Figure 1D). One of those two stable states is a focus for which the GPe-p is in a high-activity regime and forces the GPe-a to a low-activity regime. The other stable state is also a focus where the GPe-a is in a high-activity regime and forces the GPe-p to a low-activity regime. These two stable equilibria are separated by a saddle-focus. Thus, we found that strong bi-directional coupling between prototypical and arkypallidal GPe populations allows for the existence of a bi-stable activity regime, where the two populations compete over a high-activity state.

**Figure 1:**
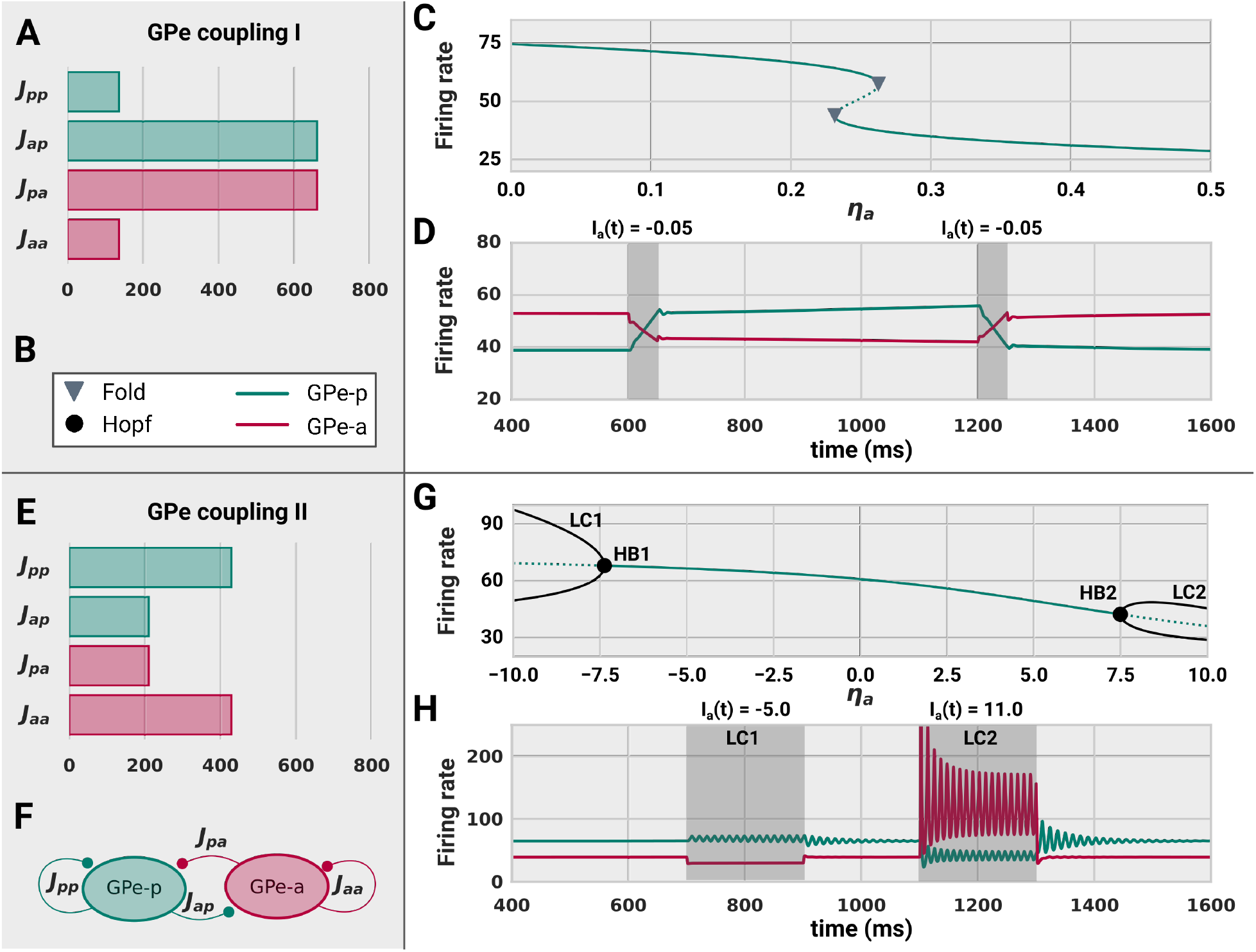
Bi-stability (**C**,**D**) vs. oscillations (**G**,**H**) in the GPe. **A:** GPe coupling strengths for *k_i_* = 2.2 and *k_p_* = 1.0. **B:** Legend of bifurcation points and line colors used in the figure. **C:** Bifurcation diagram for a 1D parameter continuation in the background input of GPe-a *η_a_*. A bi-stable regime emerges from 2 fold bifurcations for GPe coupling I and *η_p_* = 4.0. Solid (dotted) lines represent stable (unstable) solution branches. **D:** Time series of GPe-p and GPe-a firing rates that shows switching between the two stable branches via transient, extrinsic inputs to GPe-a. **E:** GPe coupling strengths for *k_i_* = 0.7 and *k_p_* = 1.0. **F:** Labels of the synaptic connections inside GPe. **G:** Bifurcation diagram for *η_a_* obtained with GPe coupling II and *η_p_* = 4.5, showing two stable limit cycles (LC1 and LC2) that emerge from two different supercritical Andronov-Hopf bifurcations (HB1 and HB2). Black, solid lines represent the minimum and maximum of the GPe-p oscillation amplitudes. **H:** Time series of GPe-p and GPe-a firing rates when forced onto the two different limit cycles via extrinsic inputs to GPe-a.

#### Coupling Pattern 2: Weak Coupling Between Prototypical and Arkypallidal Cells Promotes Oscillations

Next, we examined the consequences of a GPe coupling pattern with weak coupling between GPe-p and GPe-a and strong coupling within the populations, by choosing *k_i_* = 0.7 and *k_p_* = 1.0 (see Figure 1E). As shown in Figure 1G, both inhibitory as well as excitatory input to GPe-a pushes the system over a super-critical Andronov-Hopf bifurcation, marking the birth of a stable limit cycle. This reflects the symmetry of the chosen coupling pattern. Since GPe-a and GPe-p express strong inhibitory self-coupling with synaptic transmission delays, excitatory drive has the potential to engage each of the two populations in oscillatory behavior. However, the coupling between GPe-a and GPe-p counteracts this tendency to oscillate, if sufficiently strong. As a consequence, additional excitatory input to GPe-a engages GPe-a into oscillatory behavior (LC2 in Figure 1G), whereas additional inhibitory input to GPe-a engages GPe-p into oscillatory behavior (LC1 in Figure 1G). The latter essentially removes the desynchronizing effect that GPe-a has on GPe-p, thus enabling GPe-p oscillations which unfold around average firing rates that are in accordance with experimental recordings of GPe firing rates in monkeys (Kita et al., 2004; Wichmann and Soares, 2006).

#### Coupling Pattern 3: Strong Prototypical Projections Allow for either Bi-Stability or Periodic Oscillations

So far, we described GPe intrinsic dynamics for cases of symmetric coupling patterns between and within GPe-p and GPe-a populations. Experimental evidence suggests, however, that prototypical GPe neurons express a larger amount of axonal collaterals and synaptic connections inside GPe than arkypallidal neurons (Mallet et al., 2012; Ketzef and Silberberg, 2020). Therefore, we again performed 1D continuations in *η_a_* for multiple values of *k_i_* ∈ [0.5, 2.0] with *k_p_* = 1.5. This choice of parameters reflects the knowledge from our previous analyses that the emergence of bi-stable or oscillatory regimes critically depends on *k_i_* and changes with the level of background input to our system. The results of this analysis are summarized in Figure 2. From Figure 2A and B, it can be seen that the chosen coupling patterns allowed to enter either the bi-stable (*k_i_* = 1.8) or the oscillatory regime (*k_i_* = 0.9). In Figure 2B, we depicted the results of the 1D continuations in *η_a_* for the two different values of *k_i_*. As can be seen, the system undergoes either two fold bifurcations, or a supercritical Andronov-Hopf bifurcation, depending on the value of *k_i_*. Again, the two fold bifurcations mark the parameter boundaries of a bi-stable regime in which two stable foci are separated by a saddle-focus, whereas the Andronov-Hopf bifurcation represents the birth of a stable, periodic limit cycle. Importantly, these bifurcation can also be induced by changes in the coupling parameters of the system. This can be seen in the lower panel of Figure 2B, where we (1) identify the Andronov-Hopf bifurcation (HB2) for *k_i_* = 0.9 by decreasing the projection strength from GPe-a to GPe-p, and (2) traverse the fold bifurcations (LP3 and LP4) for *k_i_* = 1.8 by increasing the projection strength from GPe-a to GPe-p.

**Figure 2:**
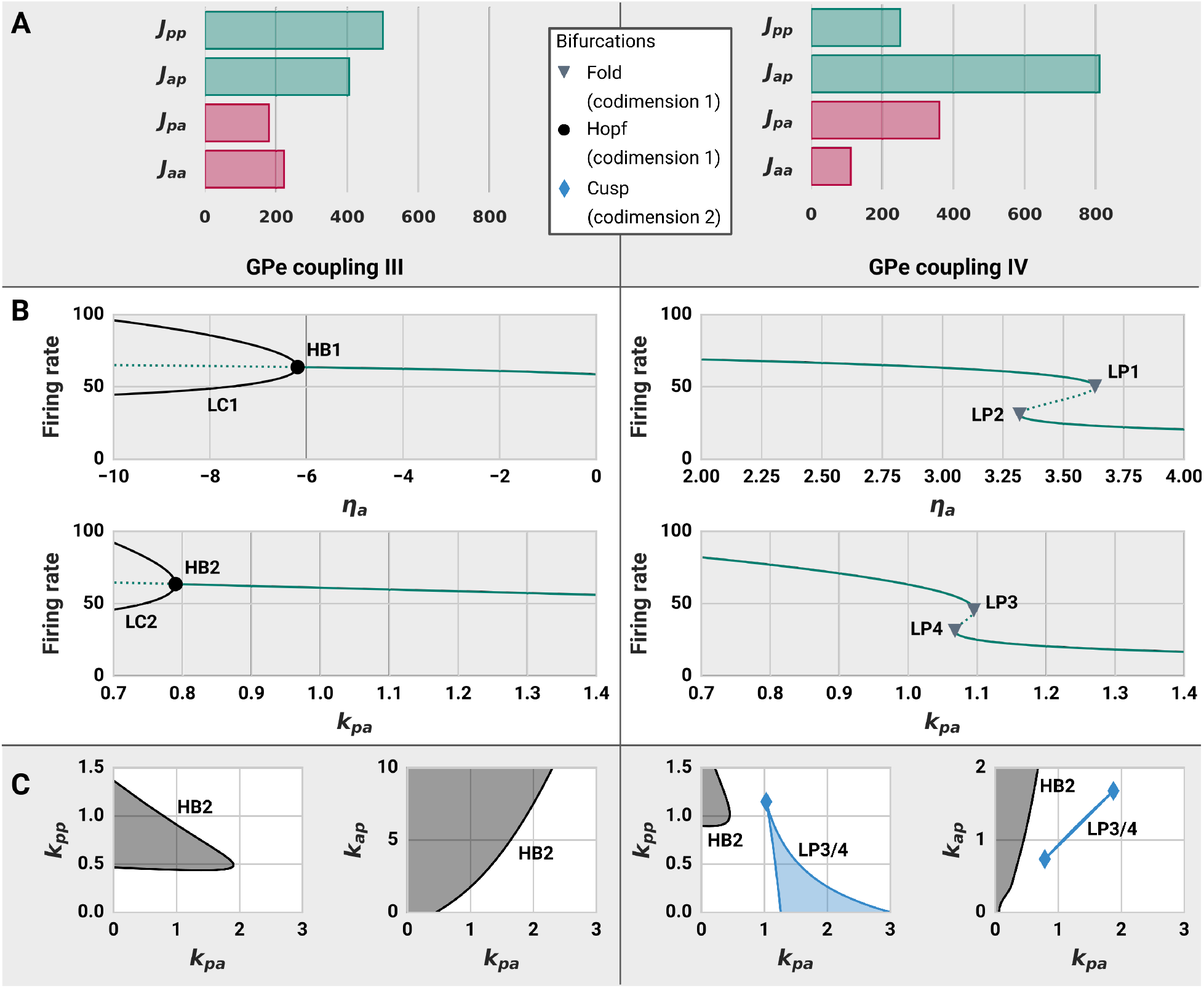
GPe phase transitions induced by changes in the synaptic coupling. **A:** GPe coupling patterns III (*k_i_* = 0.9, *k_p_* = 1.5) and IV (*k_i_* = 1.8, *k_p_* = 1.5) and a legend of all bifurcations reported in this figure. **B:** Bifurcation diagrams for 1D parameter continuations in *η_a_* and *k_pa_*. Solid lines represent stable solution branches whereas dotted lines represent unstable solution branches. For GPe coupling III and *η_p_* = 4.8, reducing *η_a_* or *k_pa_* moves the system over a supercritical Andronov-Hopf bifurcation (HB1) from which a stable limit cycle emerges. The black, solid curves show the minimum and maximum limit cycle amplitudes of the GPe firing rate. For GPe coupling IV and *η_p_* = 3.2, two fold bifurcations are found in both *η_a_* and *k_pa_*, allowing for the co-existence of two stable focus branches in a small parameter regime. **C:** 2D parameter continuations of the Hopf and fold bifurcations in the *k_pp_ − k_pa_* and *k_pp_ − k_ap_* planes for both GPe coupling patterns. Areas shaded in dark grey represent parameter regimes in which stable oscillations exist, whereas areas shaded in blue represent areas in which two stable focus branches can co-exist (bi-stability).

To test to which other intra-pallidal coupling patterns this finding generalizes, we performed pairwise two-parameter continuations in *k_pa_*, *k_pp_*, and *k_ap_*, the results of which are shown in Figure 2C. The grey-shaded areas represent areas in the 2D parameter spaces for which the oscillatory regime exists. Its boundaries are the curves of the HB2 bifurcation in the respective 2D parameter space. The Hopf curve in the *k_pp_* − *k_pa_* plane reveals that the oscillatory regime critically depends on *k_pp_* > 0, but not on *k_pa_* > 0. This confirms our earlier interpretation that this Hopf bifurcation represents an interaction of a stationary excitatory drive and a delayed, inhibitory feedback of GPe-p. Therefore, removing the inhibitory influence of GPe-a from GPe-p can drive the system over the Hopf bifurcation. Consequently, it can be seen from the *k_ap_* − *k_pa_* plane that the emergence of oscillations does not require any synaptic interaction between GPe-p and GPe-a. On the contrary, strong input from GPe-a to GPe-p prevents the emergence of oscillations.

The blue-shaded areas in Fig. 2D represent areas for which the bi-stable regime exists, and its boundaries are the curves of the LP3 and LP4 bifurcations in the 2D parameter spaces. In the two-parameter continuations of the fold curves, we identified cusp bifurcations, marking the birth of the bi-stable regime in 2D parameter space. In contrast to the oscillatory regime, the bi-stable regime does not require *k_pp_* > 0. On the contrary, the closer *k_pp_* is to zero, the broader the bi-stable regime is. Furthermore, we find that the Hopf curve can also be found for *k_i_* = 1.8, if *k_pp_* is sufficiently strong. In other words, the stable focus branching off from LP3 can undergo a supercritical Andronov-Hopf bifurcation when the synaptic input from GPe-a to GPe-p is further reduced.

### GPe Response to Periodic Forcing

By now, we have established an understanding of the intrinsic, coupling-dependent GPe response to static, afferent inputs. We found that a dichotomous organization of the GPe with two distinct populations GPe-a and GPe-p results in coupling-dependent dynamic behavior that situates the GPe either near a bi-stable or near an oscillatory regime (GPe coupling pattern III vs. IV). These two different scenarios may have substantially different consequences for the transmission and amplification of periodic input arriving at the GPe. Hence, as a next step, we analyzed the response of the GPe to periodic inputs when initialized either close to the bi-stable regime, close to the oscillatory regime, or in the oscillatory regime. To this end, we applied periodic input with period *ω* and amplitude *α* to the arkypallidal population. The input was generated by applying a sigmoidal transformation to the oscillatory signal generated by a Stuart-Landau oscillator (see equations (17)–(20) in the methods section). In each regime, we performed numerical simulations of the model behavior for different values of *ω* and *α*. We then evaluated the average phase-amplitude coupling (PAC) between the phase of low frequency signal components (2-30 Hz) and the amplitude of high frequency signal components (50-250 Hz) of the GPe-p firing rate dynamics. Furthermore, we evaluated the PPC, i.e. the phase dependency of the high frequency components on the phase of the dominating low frequency component. A detailed description of these measures is provided in the methods section. As can be seen in Figure 3, we find that the GPe responds differently to periodic input depending on its dynamic regime.

**Figure 3:**
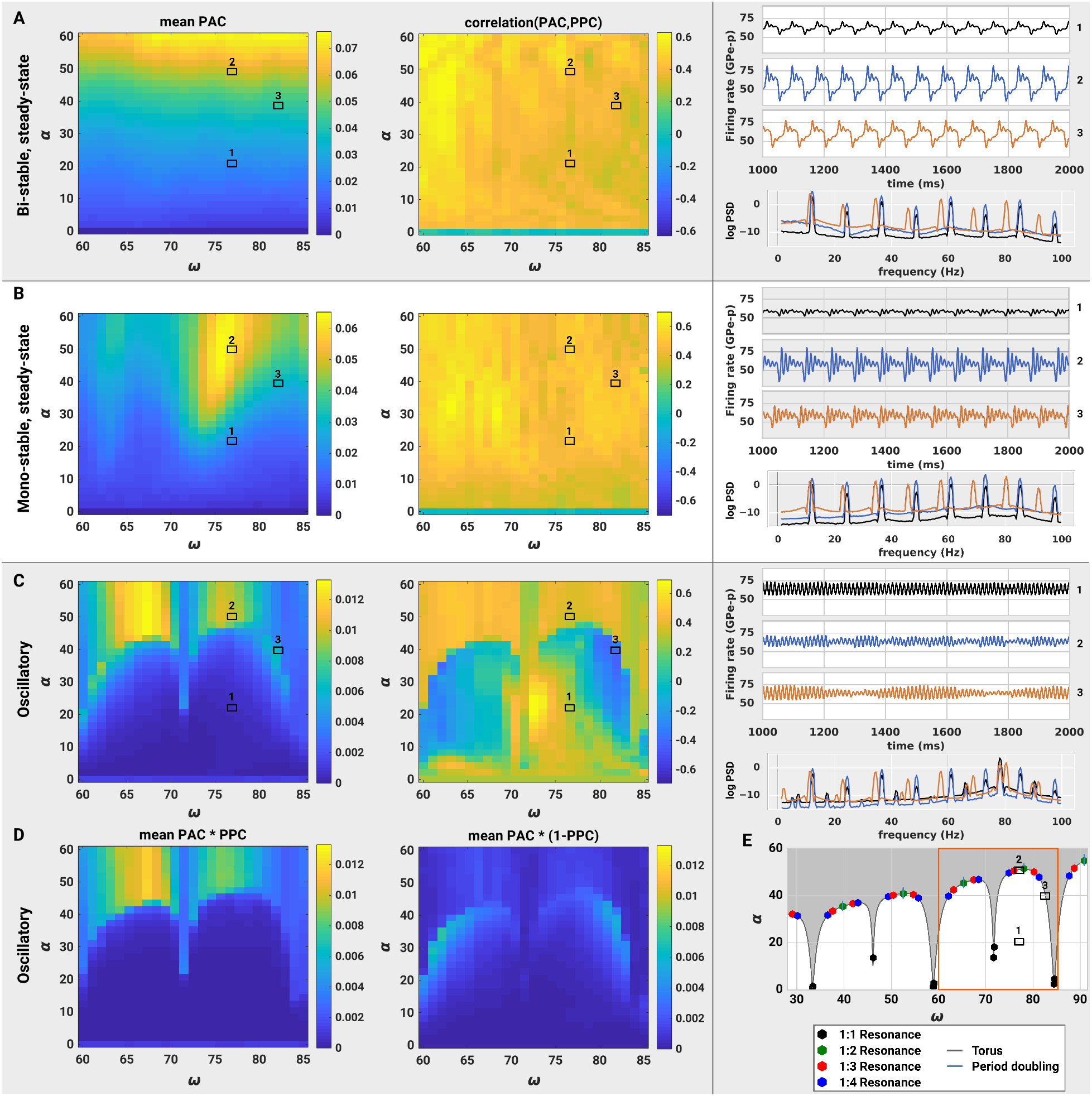
Cross-Frequency coupling between low and high frequency components in the GPe under periodic input. The input *I_a_*(*t*) was applied to the GPe-a with different frequencies 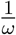 and amplitudes *α*. For each input configuration, the average phase-amplitude coupling (mean PAC) between phases of low-frequency components (2-30 Hz) and amplitudes of high-frequency components (50-250 Hz) of the GPe-p firing rate was calculated. Furthermore, the correlation between PAC and phase-phase coupling (PPC) values was calculated across all pairs of low- and high-frequency components. Exemplary time-series and log power spectral densities (PSD) are provided for the GPe-p firing rate of 3 different input configurations: (1) *ω* = 77 ms, *α* = 20, (2) *ω* = 77 ms, *α* = 50, (3) *ω* = 82 ms, *α* = 40. **A:** Results for *k_i_* = 1.8, *k_p_* = 1.5, *η_p_* = 3.2 and *η_a_* = 3.0. **B:** Results for *k_i_* = 0.9, *k_p_* = 1.5, *η_p_* = 4.8 and *η_a_* = 0.0. **C:** Results for *k_i_* = 0.9, *k_p_* = 1.5, *η_p_* = 4.8 and *η_a_* = −6.5. **D:** Product of PAC and PPC values, averaged across all pairs of low- and high-frequency components. The left (right) panel shows input configurations for which high PAC values coincide with high (low) PPC values. **E:** 2D Bifurcation diagram in the *α* − *ω* plane which shows emergence of resonant behavior and period doubling of GPe oscillations along a torus bifurcation curve.

In Figure 3A, the GPe response to periodic input is depicted for GPe coupling pattern III. Whereas small amplitude input merely perturbs the system around a stable focus, sufficiently strong input can move the system across the fold bifurcations and thus force the system in and out of the attracting domains of two different foci. This switching behavior, in combination with the relaxation to the foci, creates oscillations with interleaved large and small amplitude oscillations. Since the switching behavior happens at the input frequency, it acts as an amplification of the input. This is also reflected in the high power spectral density (PSD) of the GPe-p firing rates that can be seen at the input frequency in Figure 3A. Furthermore, the dependency of the switching on the input strength can be observed in the cross-frequency coupling. Stronger inputs generate stronger modulation of high frequency amplitudes by low frequency phases, as evaluated by PAC. Such increases in PAC occur together with increased phase locking between low- and high-frequency components. This can be observed by the generally high PAC-PPC correlations in Figure 3A.

Figure 3B shows the GPe response to periodic input when initiated close to, but not in the oscillatory regime. In this case, a stable focus is the only equilibrium, and the input perturbs the system around that equilibrium at the input frequency. After a perturbation, the system relaxes back to the focus via damped oscillations. However, since there exists no second stable equilibrium in the vicinity of this regime, the GPe is not attracted by another equilibrium when pushed away from the focus. Thus, PSDs of the GPe-p firing rates are generally shifted to higher frequencies in comparison to the bi-stable regime (see Figure 3B vs. A). Cross-frequency coupling between the phase of the stimulation and the amplitude of the focus dynamics can still be prominent though. As can be seen in Figure 3B, the PAC profile expresses the shape of a tongue, centered around *ω* ≈ 77.0ms. This tongue corresponds to a region in the *α* − *ω* space, in which the system has just enough time between subsequent stimuli to relax back to its steady state before the next perturbation pushes it away from the equilibrium state again. As a consequence, high PAC values tend to coincide with high PPC values, since a potential interference between focus frequency and input frequency cannot have a strong impact on the system dynamics.

More complex, resonant behavior can arise for periodic forcing of the GPe, if the GPe already expresses oscillations autonomously (see Figure 3C-E). When increasing *α*, the system undergoes a torus bifurcation that emerges from the interaction between the intrinsic limit cycle and the extrinsic, periodic input. As can be seen from the time series in Figure 3C, this torus bifurcation separates a regime with small, aperiodic amplitude modulations, from a regime with strong, periodic or quasi-periodic modulations of the intrinsic limit cycle. A continuation of the torus bifurcation in the *ω*-*α* plane reveals that the system expresses resonant behavior at various integer multiples of the input period (see Figure 3E). Close to regimes of 1:2 resonances, we were able to identify small loci of period doubling bifurcations, suggesting the existence of chaotic regimes. Notably, the resonant behavior could only be observed for sufficiently strong dynamic interactions between GPe-a and GPe-p. For *k_pa_* = 0.0 and *k_pp_* = 1.2 (which lies within the oscillatory regime in Fig. 2C), we identified the same torus curve as in Figure 3C, but not the resonance or period doubling bifurcations. Thus, even though the GPe-a is not required for the generation of oscillations inside GPe, it can have a substantial impact on the GPe response to periodic input. This impact is also reflected in the PAC and PPC profiles, as we find a strong dependence of these profiles on the bifurcation structure of the system. PAC values are low before the system undergoes the torus bifurcation and strongly increase beyond the torus bifurcation. In the vicinity of the torus bifurcation, we find regimes where strong PAC can co-exist with low PPC values. These regions express negative correlations between PAC and PPC and are clearly separated from regions where increased PAC and PPC co-exist (see Figure 3C). A reason for these negative correlations can be seen in the PSDs of the exemplary time series in Figure 3C. Peaks in the PSD profile correspond to harmonics of (a) the intrinsic gamma frequency of the GPe, and (b) the extrinsic beta frequency of the input. If a harmonic of (b) is in close proximity of the intrinsic gamma frequency, such as in the case of time series 3 with an input frequency of 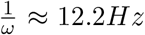, this can lead to strong resonances. The corresponding amplitude modulations are strongest at the intrinsic gamma frequency, whereas phase coupling is strongest at a harmonic of the input frequency.

### Model Generalization to GPe Spiking Neural Networks

In this section, we report how the above described findings generalize to spiking neural networks (SNNs) of coupled GPe-p and GPe-a cells with realistic cell counts and coupling probabilities. To this end, we attempted to replicate the mean-field model dynamics shown in time series 3 of Fig. 3A and C in SNNs with different network sizes and (b) different coupling probabilities. We created a total of four SNNs with (a) either all-to-all coupling or only 5 % of all possible connections, and (b) either *N_p_* = 4000 (*N_a_* = 2000) GPe-p (GPe-a) cells or *N_p_* = 40000 (*N_a_* = 20000) GPe-p (GPe-a) cells. We then repeated our simulations of the GPe response to periodic stimulation with amplitude *α* = 40.0 and period *ω* = 82.0 ms for spiking neural networks initialized near the bi-stable and in the oscillatory regime (same parameterizations as reported in Fig. 3 for the mean-field model). The dynamics of all four SNNs can be seen in comparison to the mean-field predictions in Fig. 4.

**Figure 4:**
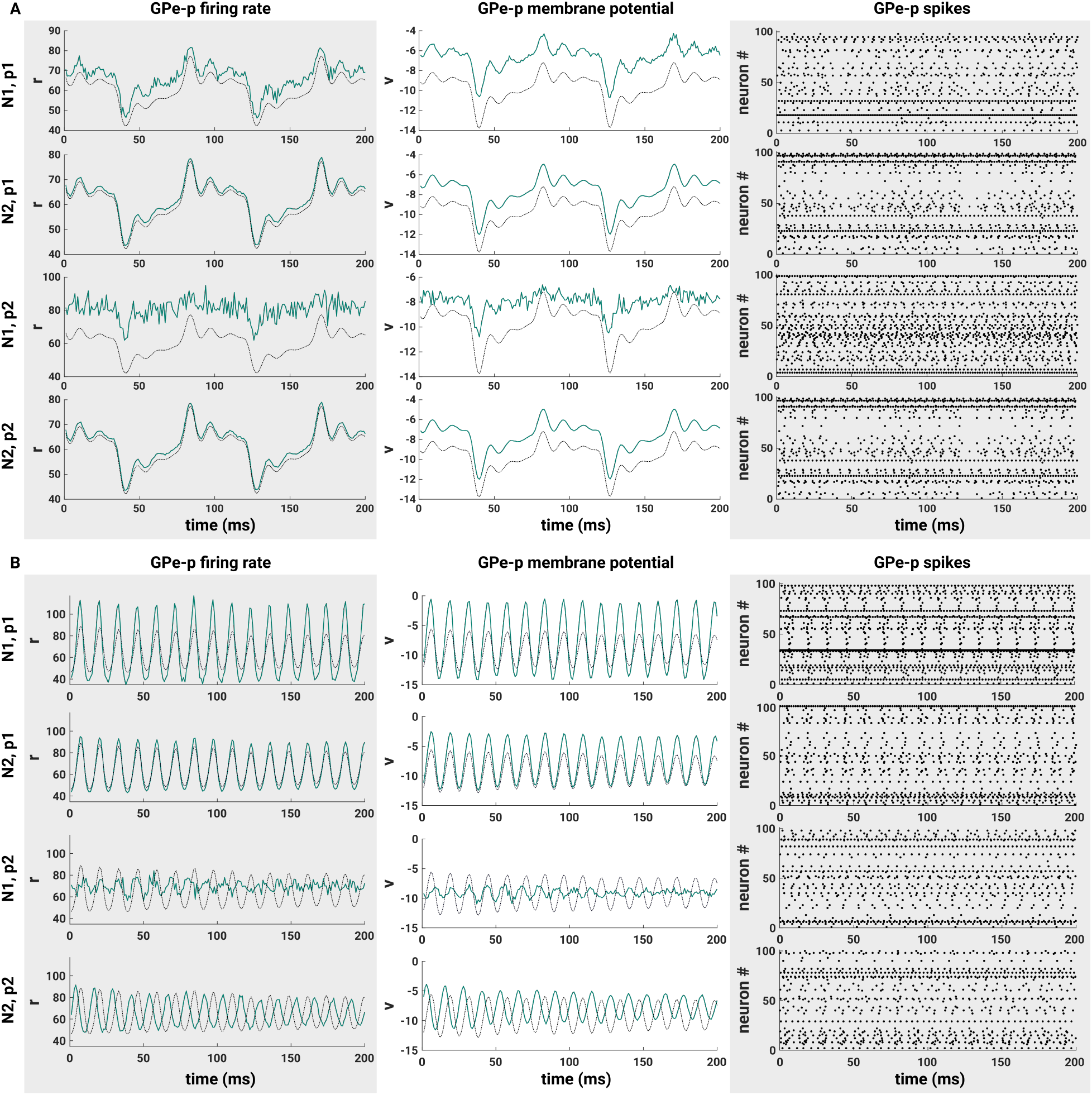
Comparison between mean-field model (black) and spiking neural networks (green). SNNs are composed of *N_p_* = 4*N*_1/2_ GPe-p and *N_a_* = 2*N*_1/2_ GPe-a neurons, where *N*_1_ = 1000 and *N*_2_ = 10000. From all possible synaptic connections in the SNN, either *p*_1_ = 100% or *p*_2_ = 5% are established. All models were driven by periodic input *I_a_*(*t*) with period *ω* = 82 ms and amplitude *α* = 40. **A:** Results for *k_i_* = 1.8, *k_p_* = 1.5, *η_p_* = 3.2 and *η_a_* = 3.0. **B:** Results for *k_i_* = 0.9, *k_p_* = 1.5, *η_p_* = 4.8 and *η_a_* = −6.5.

As expected, we find that an all-to-all coupled SNN of large size behaves nearly identical to the mean-field prediction, where the remaining difference in the oscillation amplitude is an effect of the network size and would vanish if we increased the network size even further (see difference between SNNs with *N*_1_ and *N*_2_). Interestingly, we find that reducing the number of synaptic connections to *p* = 5% of all possible connections attenuates synchronized oscillations in the network for small network sizes. However, for a sufficiently large network, the SNN follows the macroscopic dynamics predicted by the mean-field model, even when *p* = 5%. This holds for both the bi-stable as well as the oscillatory regime.

## Discussion

In this work, we investigated the implications of a dichotomous GPe structure on its intrinsic dynamic regimes and on its response to oscillatory input. This was motivated by experimental results that strongly suggest that there exist two neuron types (GPe-p and GPe-a) inside GPe with different projection targets and electrophysiological features (Mallet et al., 2012; Abdi et al., 2015; Hernández et al., 2015; Hegeman et al., 2016). Our investigations were based on populations of coupled QIF neurons, for which we derived exact mean-field equations describing the low-dimensional dynamics of average membrane potentials and firing rates. In the vicinity of realistic steady-state firing rates (*r_p_* ≈ 60*Hz* and *r _a_* ≈ 30*Hz*) (Kita et al., 2004; Wichmann and Soares, 2006; Mallet et al., 2012), we found two different phase transitions.

When we initialized the model with strong bidirectional coupling between GPe-p and GPe-a and relatively weak self-inhibition within GPe-p and GPe-a, we identified a bi-stable regime. This regime provides a form of network memory and a reliable way to switch GPe output between the different projection targets of GPe-p (STN) and GPe-a (STR). Furthermore, if the GPe is situated close to the bi-stable regime, periodic inputs from STN and STR could be strongly amplified due to a periodic switching of GPe activity between the two stable states. Indeed, by applying periodic input to the GPe-a, we were able to elicit such periodic switching in the beta frequency range, which is characteristic for PD (Brown, 2003; Hammond et al., 2007). Importantly, this also led to strong phase-amplitude as well as phase-phase coupling between beta and gamma components of the GPe-p firing rate dynamics. Interestingly, the synchronized neural activity that has been detected in recordings of STN and GPe activity from PD patients expressed not only increased power in the beta frequency band, but also increased PAC between the phase of beta components and the amplitude of gamma components (López-Azcárate et al., 2010). Assuming that GPe-a and GPe-p express strong mutual inhibition compared to their intrinsic self-inhibition, our model can explain these findings as follows: The GPe is naturally situated in the vicinity of a bi-stable regime and intrinsic changes due to PD can move the system closer to this regime. This could for example be achieved by an increased strength of the coupling between GPe-p and GPe-a. Indeed, increased synaptic efficacies have been reported in the GPe in brain slices of a rat PD model (Miguelez et al., 2012). Alternatively, increased inhibition of the GPe-p via afferent signals could also move the system closer to the bi-stable regime. A prime target for such increased afferent inhibition would be STR projection neurons, which have been demonstrated to express increased firing rates in PD (Mallet et al., 2006; Kita and Kita, 2011; Hernandez, 2014; Singh et al., 2016). By moving closer to the boundaries of the bi-stable regime, oscillatory inputs from STN or STR become more likely to elicit switching between the two stable states of the GPe. At both input sites, increased beta oscillations have been reported in PD (Brown, 2003; Belluscio et al., 2014). The switching between the two stable foci would then amplify the beta component of the oscillatory input and cause oscillatory dynamics in a gamma frequency range, thus leading to PAC. Hence, in this scenario, PD-related intrinsic changes can cause increased susceptibility of the GPe to periodic inputs, but not autonomous GPe oscillations.

When we investigated the GPe dynamics for weaker mutual inhibition between GPe-p and GPe-a, we identified oscillatory regimes near steady-state behavior. Most interestingly, inhibition of the GPe-a engaged the system in oscillations, driven by the dynamic interactions between the pace-making properties of the GPe-p and its delayed self-inhibition. Most likely, these oscillations reflect the same synchronization mechanism as reported for a single population with delayed self-inhibition in (Luccioli et al., 2019). According to their results, oscillations are counteracted by neural heterogeneity. This way, our results can be linked to the considerations in (Wilson, 2013), which suggest that strong firing rate heterogeneity together with recurrent inhibition inside GPe may serve to desynchronize GPe activity under healthy conditions. In accordance with experimental data, we modeled the GPe-a with highly heterogeneous single cell firing rates (Miguelez et al., 2012; Hernández et al., 2015; Ketzef and Silberberg, 2020). This way, inhibitory feedback from the GPe-a provides the means to suppress synchronized oscillations inside the GPe, which supports these considerations. Experimental evidence from animal models of PD suggest that GPe activity shows increased synchronization in PD (Wichmann and Soares, 2006; Mallet et al., 2012). Our model can explain these findings as follows: Increased inhibition of GPe-a removes its desynchronizing effect on the GPe-p, leading to synchronized neural oscillations within the GPe-p. Potential causes of such increased inhibition could be an increased firing rate of STR direct pathway neurons, an increased GPe-a to GPe-a projection strength, or an increased GPe-p to GPe-a projection strength. Experimental evidence suggests that each of those GPe-a inhibition mechanisms may be present in PD (Mallet et al., 2006; Kita and Kita, 2011; Miguelez et al., 2012; Hernandez, 2014; Singh et al., 2016). It has to be emphasized, though, that the emerging limit cycle leads to narrow-band gamma oscillations and thus cannot explain the emergence of beta oscillations in the parkinsonian BG (Brown, 2003; Hammond et al., 2007). However, assuming that burst-like afferent inputs drive the GPe at a beta frequency, our findings predict that GPe-intrinsic gamma oscillations can resonate with the input, leading to a waxing-and-waning of the gamma oscillations. Such waxing-and-waning behavior also implicates increased PAC, which could occur together with decreased PPC in our simulations. This finding was unique for the oscillatory regime of the GPe. Similarly, complex patterns of cross-frequency coupling have been reported previously in an instantaneously coupled two-population QIF model with sinusoidal forcing in the alpha frequency range (10 Hz) (Ceni et al., 2020). Thus, our results show under which conditions the GPe system can express the characteristic dynamics that have been identified in more abstract models of two populations with mutual inhibition.

In both GPe regimes described above, the GPe-a has a desynchronizing effect on GPe activity and can be important for explaining GPe phase transitions. Of course, the impact of the GPe-a on the overall GPe activity is limited by its coupling to other GPe cells. In principle, both GPe-p and GPe-a have been shown to express local axon collaterals within GPe (Mallet et al., 2012). A most recent study of synaptic coupling in the mouse GPe suggests that prototypical cells express considerably more GPe projections than arkypallidal cells (Ketzef and Silberberg, 2020). In our model, this corresponds to regimes where *k_p_* > 1. Furthermore, this study found that GPe-intrinsic projections of GPe-p cells preferentially target GPe-a cells over other GPe-p cells, which corresponds to *k_ap_* > *k_pp_* in our model. Finally, the study found that Gpe-a stimulation failed to inhibit both GPe-p and GPe-a cells significantly (Ketzef and Silberberg, 2 020). While this could indicate that GPe-a cells express little projections to other GPe cells, our results suggest that this finding can also be explained by a GPe regime with high GPe-p and low GPe-a steady-state activity. In such regimes, excitatory input to GPe-a had very little effect on average GPe firing rates in our model, even though GPe-a neurons expressed substantial projections to other GPe cells. Furthermore, our SNN simulations revealed that the effect of increased GPe-a firing on GPe-p neurons depends on the intrinsic firing rate of the latter. Strong pacemaker cells were barely affected, while cells with low intrinsic firing rates expressed visible firing pauses in response to GPe-a stimulation. Thus, our results can explain why (Ketzef and Silberberg, 2020) did not find a significant decrease of GPe-p firing rates upon GPe-a stimulation, even though (Mallet et al., 2012) found indicators of GPe-a to GPe-p axons. Importantly, GPe-a cells may still have a desynchronizing effect on the overall GPe activity in such a regime, by acting on spike timings rather than firing rates. This is supported by particularly heterogeneous GPe-a firing rates (Miguelez et al., 2012; Hernández et al., 2015; Ketzef and Silberberg, 2020), such that even relatively sparse GPe-a axon collaterals could serve as an important desynchronization mechanism (Wilson, 2013). If future investigations should find that GPe-a to GPe-p projections are negligible, our results predict that a GPe-p dominated focus governs GPe behavior, which may turn into a limit cycle under pathological conditions, such as increased GPe-p to GPe-p coupling (Miguelez et al., 2012). The latter scenario does not rely on the existence of a GPe-a to GPe-p connection and the role of the GPe-a would be restricted to its feedback to STR. Since the GPe-a seems to integrate information from GPe-p, STN and both striatal pathways, however, we argue that it should in any case be included in future models of BG interactions (Ketzef and Silberberg, 2020).

The mathematical model presented in this paper can serve as a basis for future BG models. We have shown that a mean-field model derived under the assumptions of all-to-all coupling and an infinite number of neurons accurately describes the dynamics of a SNN with realistic cell counts and coupling densities (Hegeman et al., 2016). Thus, our model lends itself to multi-scale approaches. It can easily be extended by additional biological details, such as plasticity mechanisms (Gast et al., 2020) or gap junctions (Pietras et al., 2019). The latter may be of particular interest to investigate parkinsonian conditions inside the GPe (Schwab et al., 2014).

## Acknowledgements

We would like to thank Bastian Pietras and Ernest Montbrió for helpful discussions. Richard Gast was funded by the Studienstiftung des deutschen Volkes. Helmut Schmidt was supported by the German Research Foundation (DFG (KN 588/7-1) awarded to Thomas R.Knösche via Priority Program 2041, “Computational Connectomics”).

